# Targeting the nervous system of the parasitic worm, *Haemonchus contortus*, with quercetin

**DOI:** 10.1101/2021.07.11.451932

**Authors:** Vanshita Goel, Neloy Kumar Chakroborty, Sunidhi Sharma, Lachhman Das Singla, Diptiman Choudhury

## Abstract

High Prevalence of infection rate, limited choice of drugs and emerging resistance against these leads to an pressing need for the development of new antihelminthic drugs and drug targets. However, limited understanding of the physiology of worms has delayed the process substantially. Here, for the first time, we are reporting the tissue morphology of *Haemonchus contortus*, and target its nervous system with a quercetin, a naturally occurring flavonoid. Quercetin showed anthelmintic activity against all developmental stages of the nematode. Histological analysis demonstrated damage of various body parts including isthmus, brut, pseudocoele, and other organs. Mechanistic studies revealed the generation of oxidative stress and alteration of the stress response enzyme activities. Importantly, the time-dependent imaging of ROS-generation revealed the neuronal system as the primary target of quercetin in the adult worms, which eventually leads to the paralysis and death of the worms. This work demonstrates neuronal system as a novel drug target for quercetin.

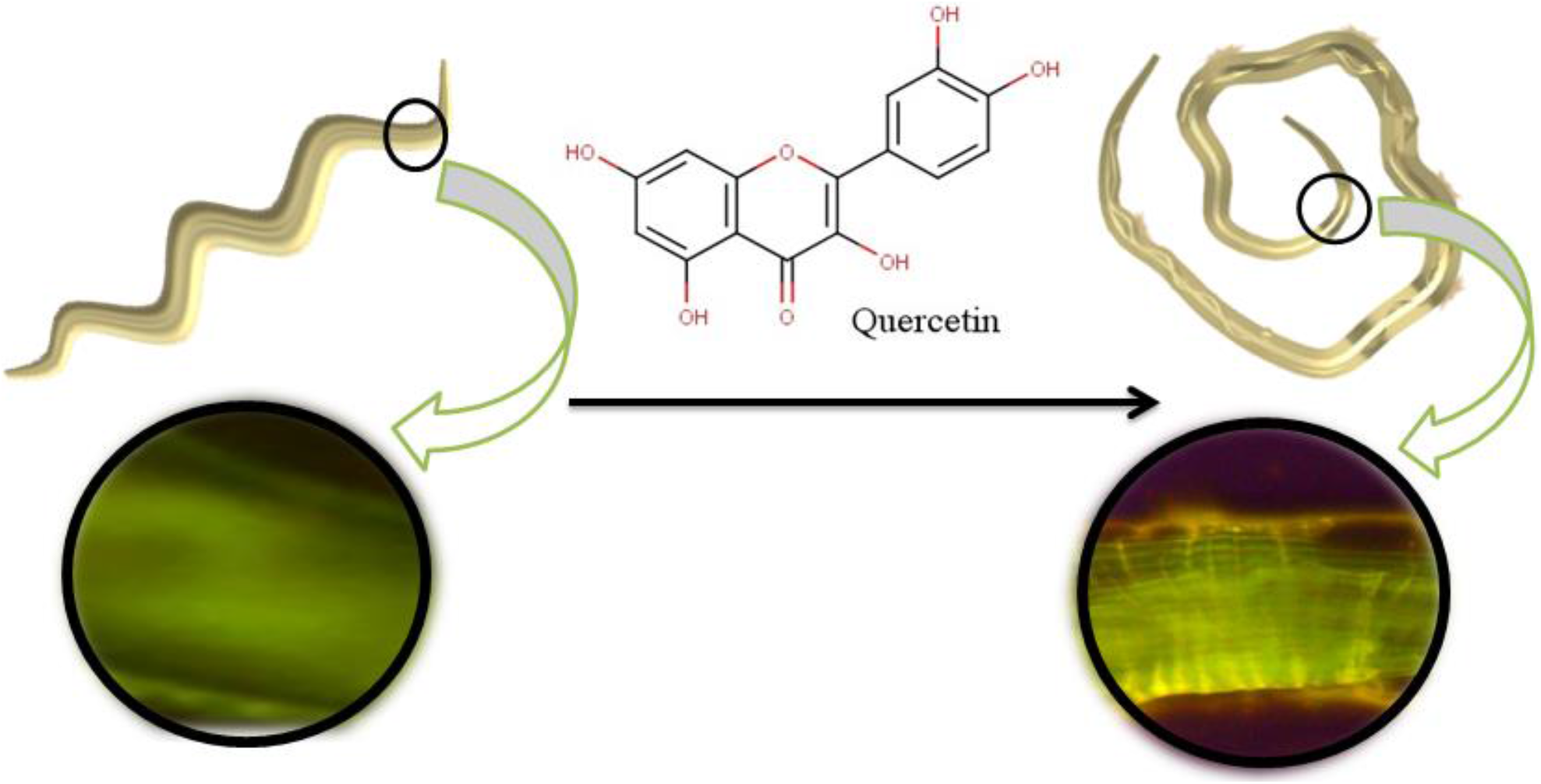
Graphical Abstract. Schematic representation showing generation of Reactive Oxygen Stress in the nervous system of *H. contortus* due to quercetin treatment.

## 1. Introduction

In the phylum Nematoda, the number of species ranges between 100,000 and 10 million, however only 20,000–30,000 species have been characterized [1]. In particular, *Caenorhabditis elegans* (*C. elegans*) is the most studied nematode, which is used as a popular genetic model system across the biological disciplines to study complex behavior, drug responses, microbiome, and pathophysiology of human diseases [2–10]. The nervous system of *C. elegans* has been studied extensively and only for *C. elegans*, the complete system-wide neural-connectome is known [11–14]. Nevertheless, the overall designs of the nervous systems in nematodes are found to be largely similar across the phylum[15]. The structure displays the invariant features of the nerve ring as the central nervous system at the anterior end of the body, the ganglia, the ventral cord that runs longitudinally throughout the body, and the commissural connections. Also, the similarity in the total number of neurons and their anatomic correspondence are remarkable [15]. These similarities have helped us immensely to understand the functioning of the neural circuits and drug-related toxicity responses in the nematodes [16–18]; of great importance is the increasing developments of drug-resistance in the intestinal parasitic helminths against the available anthelmintic drugs, such as benzimidazoles, levamisole, pyrantel, avermectins, etc [19– 27].

Among the intestinal parasitic helminths, *H*. ***c**ontortus* is a highly pathogenic candidate, causing gastrointestinal infections, primarily and not restricted to the small ruminants, with global distribution [28,29]. Due to its blood-feeding behavior and the potential for the rapid development of large burdens, it is a frequent cause of mortalities in sheep, cattle, goats, and other ruminants as well as is the most important parasite of the livestock in the warm climatic regions, and arguably on a global basis [30,31]. *H. contortus* causes anemia and infirmity, which further is responsible for severe economic losses in sheep and goat breeding globally [32]. Importantly, multiple reports of infections from different continents, with *H. contortus*, has raised serious veterinary concerns while considering the development of resistance in this parasite against the available anthelmintic drugs [32–36]. Anthelmintic resistance is now a widespread problem, especially in *H. contortus*, due to its enormous genetic diversity, which possibly is allowing the anthelminthic resistance alleles to be rapidly selected [37–39]. In 1983, Cawthorne and Whitehead have reported the development of resistance against benzimidazole in *H. contortus*[40]. Other studies showed the mechanism of benzimidazole resistance due to point mutations in the β-tubulin gene; the target site of the drug [41–44]. *In vitro* development of resistance in *H. contortus* against thiabendazole also has highlighted the significance of point mutation (from glutamate to alanine) at codon 198 of the β-tubulin gene which is expressed *in vivo* relevance [45,46]. *H. placei*, which is one of the other most threatening parasitic nematodes of cattle in the tropical areas, also developed molecular resistance mechanisms against ivermectin, which includes the efflux of the drug from the cells and the eventual ineffectiveness of drug action [47–49].

To cope up with the worldwide problem of the developments of resistance in *H. contortus* against the anthelmintic drugs, researchers are actively engaged in discovering alternative plant-based anthelmintics [50]. Screening of phytochemicals with anthelmintic properties against *H. contortus* although has identified multiple candidates, however, their use as potential therapeutics can be hindered due to lack of knowledge on their mechanisms of action [51–55]. Thus a great demand exists for better anthelmintic remedies which can target specific system to cause quick mortality in the parasitic worms. To this end, we have designed our study to investigate the anthelmintic activity of quercetin, a phytochemical, against *H. contortus*. The choice of Quercetin was particularly based on its therapeutic properties[56]. Quercetin is a naturally occurring flavonoid, present in onions, green tea, apples, berries, garlic, etc [57] and is well recognized for its anti-cancer, anti-inflammatory, and antioxidant properties [58–61]. Besides, quercetin has multiple intracellular molecular targets with the potential to reverse treatment resistance [62]. In a recent publication, it has been shown that quercetin in combination with ivermectin is capable of causing mortality in the ivermectin-resistant larvae of *H. contortus* [63-64]. Based on our *in vitro* observation in the present work, we found, for the first time, that quercetin, acts as an effective anthelmintic remedy and targets the nervous system of the parasitic nematode, *H. contortus*. Our investigation also demonstrated, for the first time, the tissue morphology and histopathology in *H. contortus* caused by the exposure to quercetin. We studied the novel mechanism of action of quercetin and found it to inflict tissue-damage and eventual death of the adult worms through oxidative-stress. The generation of reactive oxygen species in the nervous system of *H. contortus* and eventual paralysis and mortality of the adult worms are extremely interesting findings keeping in mind that quercetin has characterized roles in the nervous system as neuroprotective (inhibits oxidative stress mediated neuronal damage), anti-neuroinflammatory (suppresses of NF-κB sinagling, clearance of β-amyloid peptide and hyperphosphorylated tau) and neurogenesis agent (increases neuronal longevity by modulating signaling pathways mediated by NGF, BDNF, and various kinases)[56]. we have discussed the results from the perspectives of repurposing an old drug, quercetin, for a novel function and its potential to be used in clinical studies as a drug target for the nervous system of the parasitic nematodes.

## 2. METHODS

### 2.1. Collection of *Haemonchus contortus* from the ruminants

The abomasa infected with adult *H. contortus* worms were isolated from the slaughterhouses of Ludhiana, Punjab, India. The abomasa were washed thoroughly with running water to expel out the contaminations. After disinfection, the abomasa were dissected and the internal contents were collected, and then washed several times using a sieve. *H. contortus* adults were singled out and identified using a fine brush and were readily transferred to 1× phosphate buffer saline (PBS) at pH 7.4[65].

### 2.2. Isolation of eggs from the adult females

The egg islation was carried out from the adult female worms using particular guidelines with some modifications[66]. Adult females were identified, based on their morphological characteristics and washed with 1× PBS to remove the contaminations. They were centrifuged at 11,000 x g for 15 min in the freshly prepared saturated sterile saline. Finally, a suspension was diluted in 1 ml of 1× PBS to obtain a concentration of ∼200 eggs/ml using the McMaster technique[67] and stored at 4 °C for further use.

### 2.3. Selection and reaping of the infective larval stage

L3 larvae of *H. contortus* were harvested from the infected fecal samples of infected ruminants after 7 days of culturing period. The mixed fecal culture was incubated for 7 days at room temperature (25-31°C). Regular supplementation of water kept the mixed culture wet for better oxygenation and humidity. Thereafter, the mixed culture was analyzed and the infective larvae were collected from the mixed fecal culture and stored at 4 °C for the anthelmintic tests. The larvae were cleaned with 0.2 % sodium hypochlorite solution to remove the adherent bacteria and then finally used for the *in vitro* larval mortality assay[68].

### 2.4. Adult paralysis and mortality assay

Mortality test for the adult *H. contortus* was conducted using 15 male and 15 female worms, in a 35 mm culture plate in presence of quercetin at different concentrations (0.125, 0.25, 0.50, 1 and 2 mM) and Albendazole (0.2 mM in 1× RPMI medium) (Roswell Park Memorial Institute-1640, Lot: 98H83171) for 24 h at room temperature (37 °C). After the incubation, the mortality of the worms, morphology, and responses to the physical stimuli was assessed directly under the microscope at 40× magnification. Paralysis and death times were calculated at regular time intervals (0, 1, 3, 6, 12, and 24 h). The mobility of adult worms was analyzed after visualizing their movement for 60 sec under moderate agitation. Further, the mortality of the treated worms was confirmed by checking their responses to the heat shock in the saturated saline solution at 50 °C for 10 secs and consequently they were stained with Lugol’s stain to observe the morphological changes[69, 70,].

### 2.5. Larval mortality assay

*In vitro* mortality test of *H. contortus* larvae was performed by taking the L3 larvae (approx 25-30) in 200 µl of RPMI medium in the 96 well plates in presence of quercetin at different concentrations (0.125, 0.25, 0.50, 1, 2, and 5 mM) and Albendazole (0.2 mM) for 24 h. After the incubation, direct microscopic examination (at 40 × magnification) was performed to check for mobility (for 20 sec) to the physical stimuli. The morphology of the larvae was also monitored. Larvae, other than the motile ones, were considered as dead. Thereafter, Lugol’s staining was performed to check the morphological changes[71]. Percentage of larval mortality (% ML3) was calculated using the formula:

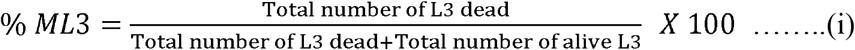

### 2.6. Egg hatching assay

The experiment was performed using 1 ml of egg suspension, containing nearly 200 eggs, into 24 well plates, and subsequently, the efficiency of quercetin to cause inhibition in egg hatching was assessed. The same set of concentrations of quercetin was used (0.125, 0.25, 0.50, 1, 2, and 5 mM) along with the controls and Albendazole (0.2 mM). Subsequently, the microtiter plate was incubated at 28 °C for 48 h. The treatments were carried out in triplicate to calculate the average percentage of inhibition in egg hatching. After 48 h of the treatments, Lugol’s iodine drop (10 µl) was added to each well to discontinue the inhibition process, and the final counts of the number of eggs hatched and inhibited were obtained under the microscope. The percentage of inhibition to egg hatching (% EHI) was calculated using the formula:

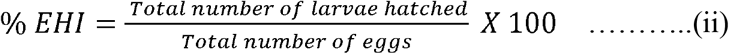

### 2.7. SEM studies for monitoring physical damage to the worms

Scanning electron microscopy was performed to monitor the physical damage in the adult worms caused by quercetin treatment. Worms were treated with 1mM of quercetin and 0.2 mM of albendazole for 12 h and then washed with 1× PBS. After the PBS wash, treated and control worms were dehydrated using increasing concentrations of ethanol from 50% to 100%. After the dehydration, samples were coated using gold (15 μm) under a specific vacuum condition. The gold coated specimens were then observed in the SEM (JSM-6490LV Scanning Electron Microscope, JEOL, USA) at an electron accelerating voltage of 15 KeV[72].

### 2.8. Histopathological investigation in *H. contortus*

Hematoxylin and Eosin (H&E) staining was done on the female worms treated with 1 mM of quercetin and Albendazole (0.2 mM) at room temperature and samples were stored in 99.9% ethanol (MERCK). Ascending dehydrations of the worm tissues were performed overnight. Wax blocks were prepared by embedding the worm tissues inside. Blocks were stored at -20°Cto to harden for sectioning. Sections were cut in the microtome (thickness of the sections was 0.2 mm) and readily transferred to 50°C hot water to remove the extra wax present on the tissue. Then the tissue was procured with a glass slide and kept on a 65 °C hot plate to allow the extra wax to melt for the fixation of the worm on the slide. Staining was done following the procedure that is described previously. The slides were kept for 2 min in different concentrations of ethanol and then 1 min each in the H and E stains. Further, microscopic examinations were done to identify the morphological changes[73].

### 2.9. Detection of reactive oxygen species

Worms were treated with quercetin (1 mM) for 3 h and thereafter washed with the distilled water. Further, the worms were treated with 100 nM of DCFDA (2’,7’-dichlorodihydrofluoresceindiacetate) and incubated for 20 min in the dark at 37°C. After incubation, samples were washed with distilled water to remove the excess of DCFDA and fixed in the slide for fluorescent imaging (fluorescence inverted microscope, Dewinter, Italy). ROS generation by quercetin was observed at the anterior and posterior ends, and body regions, and was compared with the Alb-treated (0.2mM) and RPMI Media-treated controls[67].

### 2.10. Measurements of antioxidant enzyme activities

Total protein concentrations in the treated and control worms were calculated using the Bradford method. For the Bradford assay, 250 mg of the treated and control helminth tissues were homogenized using 500 µl of RIPA buffer (Radioimmunoprecipitation assay). The total protein concentration was estimated in the clear supernatant of the homogenized mixture, after centrifugation at 10,000 rpm for 15 min, from the BSA standard curve. The superoxide dismutase (SOD) activity was determined by the spectrophotometric method of Kong et al.[74]; Catalase activity (CAT) was determined following the method of Hadwan and Abed[75]; and GSH-Px with the method reported by Antunes and Cadenas[76]. Total enzyme activity was determined by using the following equation (iii).

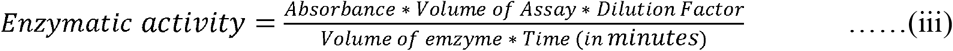

### 2.11. Statistical Analysis

Data were presented as the means and standard deviations of the independent experiments (experiments were triplicated). LD_50_ values for adult mortality, larval mortality, and egg hatching were calculated using probit analysis. To understand the dependence of larval mortality, egg hatching, and enzymatic activities on the concentration of quercetin, we performed oneway ANOVA test. Following the one-way ANOVA, Tukey’s HSD posthoc test was conducted to compare the means of the different quercetin-treated groups (different concentrations of quercetin) for these three parameters. T-test for independent samples was performed to compare between the means; control vs. Alb-treated groups and between the Alb-treated and quercetin group. Repeated measures ANOVA followed by Tukey’s HSD posthoc test were performed to check whether generation of ROS, due to quercetin-treatment, was increasing with time. For this, we selected a 10×10 pixel area within the nerve ring portions of the images (captured at multiple time points) and used the pixel intensity values for the repeated measures ANOVA test. Two measurements were considered statistically significant if the corresponding p-value was <0.01. Statistical analysis of data was conducted by using SPSS and images were prepared using Matlab and Microsoft PowerPoint.

## 3. Results

### 3.1. Adult paralysis and mortality assay

In the male adult worms, quercetin was found to be most active at the concentration of 1 mM, at which 40 % of the worms were paralyzed after 1 h and 80 % were paralyzed after 3 h of exposure. At 6 h, 40 % were dead and after 12 h, we found 80 % mortality in the adult worms (Table 1). Furthermore, in adult males, quercetin at 1mM concentration caused 100 % mortality within 24h after the treatment. In the case of adult females, quercetin showed a slower effect than the males as 40 % paralysis of the adults took place after 3 h and only at 6 h, 80% of the worms were paralyzed. Mortality in the females was only visible at 12 h (60 %) however, 100 % mortality was found after 24 h of quercetin treatment (Table 1). The LD_50_ values for adult mortality calculated respectively for the adult male and female worms were 0.62 mM (95 % lower confidence limit 0.49, 95 % higher confidence limit 0.78) and 0.88 mM (95 % lower confidence limit 0.71, 95 % higher confidence limit 1.1) at 12 h of quercetin exposure. Physical damage under the bright field has also been demonstrated in the control, Alb (0.2 mM) and Quercetin (1 mM) treated worms respectively (Fig. S1 a-c).

**Table 1:** Death time and Paralysis time analysis of adult male and female *H. contortus* due to quercetin treatment.

| Time of Exposure | Paralysis Time |  |  |  |  |  |  | Death Time |  |  |  |  |  |  |
| --- | --- | --- | --- | --- | --- | --- | --- | --- | --- | --- | --- | --- | --- | --- |
|  | Male Worms |  |  |  |  |  |  |  |  |  |  |  |  |  |
|  | Con | Alb | 0.125 | 0.25 | 0.5 | 1 | 2 | Con | Alb | 0.125 | 0.25 | 0.5 | 1 | 2 |
| 0 hr | - | - | - | - | - | - | - | - | - | - | - | - | - | - |
| 0.5 hr | - | - | - | - | - | - | 9±0 | - | - | - | - | - | - | - |
| 1 hr | - | - | - | - | - | 6±0 | 12±0.5 | - | - | - | - | - | - | - |
| 3 hr | - | - | - | - | 9±0 | 12±0.5 | 3±0.5 | - | - | - | - | - | - | 12±0.5 |
| 6 hr | - | 3±0.5 | - | - | 12±0.5 | 9±0 | 1.5±1.0 | - | - | - | - | 3±0.5 | 6±0 | 13.5±1 |
| 12 hr | - | 7.±0.5 | 3±0 | 6±0.5 | 9±0 | 3±0.5 | 0 | - | 1.5±1 | - | - | 6±0.5 | 12±0.5 | 15±0 |
| 24 hr | 1±0 | 0 | 9±0.5 | 12±0.5 | 0 | 0±0 | 0 | - | 15±0 | - | 3±0 | 15±0 | 15±0 | 15±0 |
| Female Worms |  |  |  |  |  |  |  |  |  |  |  |  |  |  |
| 0 hr | - | - | - | - | - | - | - | - | - | - | - | - | - | - |
| 0.5 hr | - | - | - | - | - | - | - | - | - | - | - | - | - | - |
| 1 hr | - | - | - | - | - | 3±0.5 | 6±0 | - | - | - | - | - | - | - |
| 3 hr | - | - | - | - | 3±0 | 6±0.5 | 12±0.5 | - | - | - | - | - | - | 3±0 |
| 6 hr | - | - | - | - | 9±0.5 | 12±0 | 6±1.5 | - | - | - | - | - | - | 9±1.5 |
| 12 hr | - | - | - | 3±0 | 15±0 | 6±0.5 | 0 | - | 3±0 | - | - | - | 9±0.5 | 15±0 |
| 24 hr | 1±0 | 9±1 | 3±0 | 3±0.5 | 4±0.7 | 0 | 0 | - | 6±1 | - | 3±0 | 11±0.7 | 15±0 | 15±0 |

### 3.2. Larval mortality assay

Spindling shrinkage and tissue damage morphology of L-3 larvae were observed in the case of quercetin treatment, whereas no such change was found in the albendazole-treated samples. The survival of larvae was measured after 24 h of quercetin treatment using equation (i). The reduction in the percentage of larval survival with the increase in quercetin concentration was seen: 60.0±6.6, 33.3±3.3, 18.8±3.8, 10.0±3.3, 1.1±1.9 and 0 % survival respectively for 0.125, 0.25, 0.50, 1, 2, and 5 mM of quercetin. In contrast, albendazole (Alb) showed 56.6±3.3 % survival of the L-3 larvae at 0.2 mM (Fig. S2). We measured an LD_50_ value of 0.16 mM (95% lower confidence limit 0.09, 95% higher confidence limit 0.22) for the larval mortality for 24 h of quercetin treatment. A one way ANOVA test showed a significant interaction between the larval survival and concentration of quercetin (F_(5, 12)_ = 111.31, *p*<0.00001), which confirmed the dose-response effect for quercetin (Results of the posthoc test, comparing larval survival for the pairs of quercetin concentration, are given in the supplementary Table ST1). Albendazole (Alb) treatment also showed significantly reduced survival of the L3 larvae compared to the control (T-test for independent samples: t = 19, df = 4, *p* = 0.00004). We also compared the survival of larvae treated with Alb with a comparable concentration of quercetin, i.e. 0.25 mM, and found significantly less survival in the quercetin treated than the Alb treated larvae (T-test for Independent Samples: t = -8.57, df = 4, *p* = 0.001).

### 3.3. Egg hatch assay

For the egg hatch assay, percentages of inhibition in egg hatching were measured after 48 h of quercetin treatment and quantified using equation (ii). The results, demonstrating the toxic effect of quercetin, showed reduced egg hatching with the increase in quercetin concentration: 69.5±2.5, 35.3±3.3, 29.5±4.2, 19.5±1.5, 15.6±2.3, and 10±1.8% of egg hatching (presence of larvae L1) respectively at the concentrations of 0.125, 0.25, 0.50, 1, 2, and 5 mM (Fig. S3). Contrastingly, albendazole (0.2 mM) showed 72.6±3.01% egg hatching. LD_50_ measured for the inhibition of egg hatching was 0.19 mM (95% lower confidence limit 0.04, 95% higher confidence limit 0.37) for 48 h. A one-way ANOVA test showing a significant interaction between the egg hatching and concentration of quercetin (F_(5, 12)_ = 177.78, *p*<0.00001) has confirmed the dose-response effect for quercetin (80 % of the eggs fail to hatch at 1 mM). Results of the posthoc test, comparing the number of eggs hatched for the pairs of quercetin concentration, are given in the supplementary Table ST2. Similar to the findings of the larval mortality assay, Alb treatment also showed significantly less egg hatching than the control (t = 7.68, df = 4, *p* = 0.001) whereas quercetin at 0.25 mM showed significantly less egg hatching compared to the Alb treatment (t = -14.39, df = 4, *p* = 0.0001).

### 3.4. Morphological damage inflicted by quercetin

Scanning electron microscopy was performed on the adult worms to understand the possible morphological changes caused by exposures to albendazole and quercetin. In the control group, treated with RPMI Media, smooth cuticles with well-developed body regions were seen in the adult female worm (Fig. 1a). In the anterior portion, intact sharp end and blood-sucking mouth region with no disruption (Fig. 1b) were identified. In the posterior end of the worm, a sharp end with an intact tail region was identified (Fig. 1c). In the case of Alb-treated (0.2 mM for 3 h) worm, folds, partial shrinkage, and disorganization of the cuticle were observed throughout the body (Fig. 1d) as well as at the anterior (Fig. 1e) and posterior (Fig. 1f) ends. However, in the quercetin-treated worm (1 mM for 3 h), complete disorganization and disruption of body regions along with the loss of the cuticle were observed (Fig. 1g). Also, shrinkage of the body ends, a complete loss of the blood-sucking anterior mouth region, and rupturing of the cuticle at the tail region were observed (Fig. 1h-i).

**Fig. 1:**
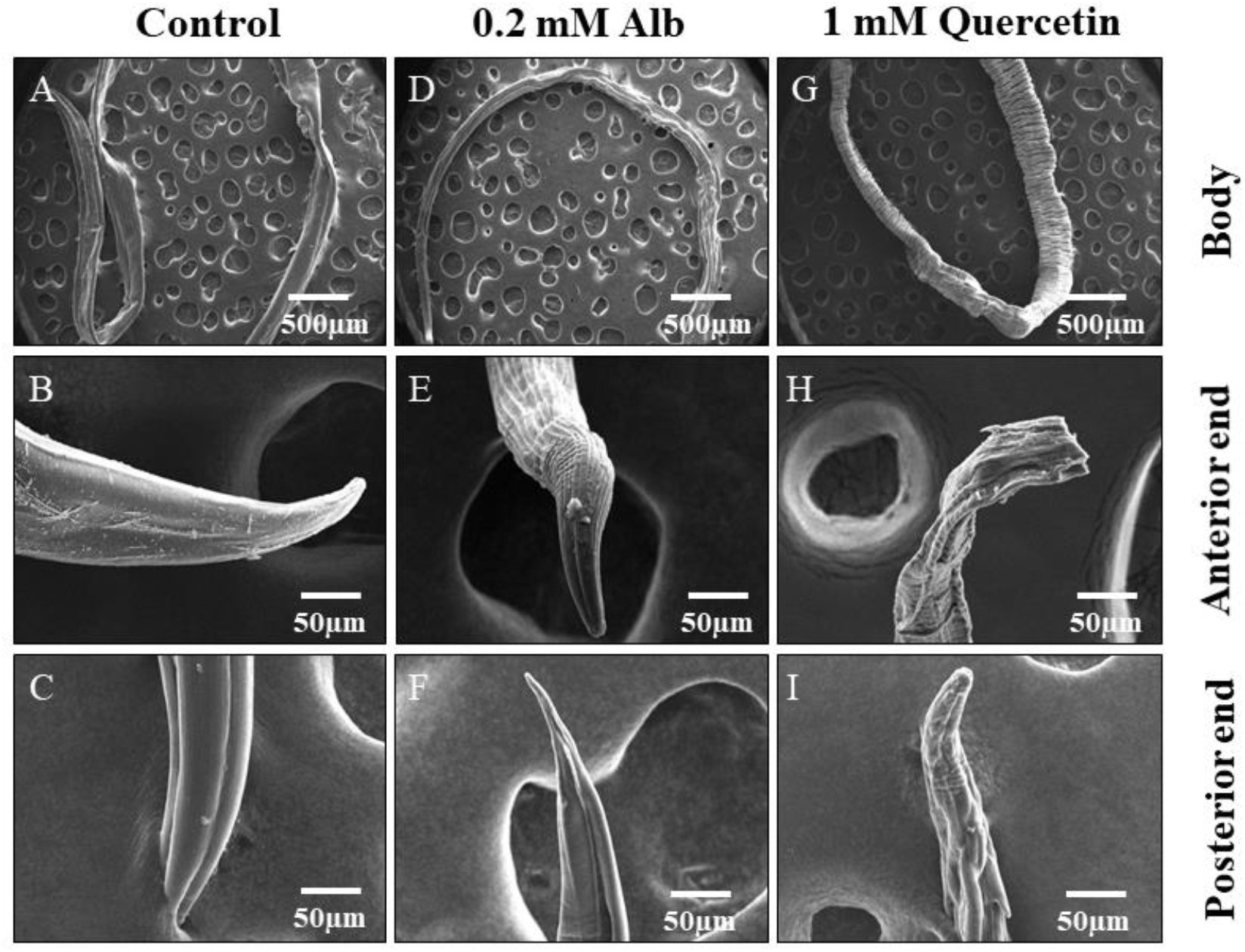
Study of morphological damage in the adult *H. contortus* due to the treatment with quercetin. Scanning electron microscopic images of the adult *H. contortus* are shown here for the three sets of conditions. The first three images (**a, b**, and **c**) are showing the intact body and the anterior and posterior ends in the control group, treated with RPMI media. The second set of three images (**d, e**, and **f**) is showing partial disruptions of the body ends and cuticle due to the treatment with 0.2 mM Alb. The last set of three images (**g, h**, and **i**) is showing the profound damage in the body structures and disruptions in the cuticular integrity due to the treatment with 1 mM of quercetin.

### 3.5. Histopathology caused by quercetin treatment

Massive changes in the morphology, caused by quercetin treatment, were observed under the light microscope (Fig. 2c) whereas Alb (Fig. 2b) showed little to no change in the morphology in comparison to the control (Fig. 2a). Detailed magnified image analysis revealed intact anterior and posterior ends with unblemished (ai) isthmus, (aii) brut, (aiii) pseudocoele, (aiv) globular leukocytes (surrounded by small-sized cells), (av) muscle cells, (avi) intestinal epithelial region, (avii) ovum and growth zone of the ovary, and (aviii) intact skin tissue in control adult worm (Fig. 2a). In the 0.2 mM Alb-treated worms (Fig. 2b), moderate disruptions were visible at the (bi) isthmus and (biii) pseudocoele along with the (bvi) partially-punctured muscle cells, (bv) globular leukocytes with less number of surrounding cells, and a (bviii) ruptured ovary. On the other hand, quercetin-treatment (Fig. 2c) has caused complete disruption of the anterior end, a near-total disruption of the ovary, and loss of eggs, splitting of the pseudocoele, punctured muscle cells, and damaged globular leukocytes with limited counts of surrounding-cells. Fig. 2c ci-cviii is showing the histopathology of different body parts in the adult female *H. contortus* treated with quercetin.

**Fig. 2:**
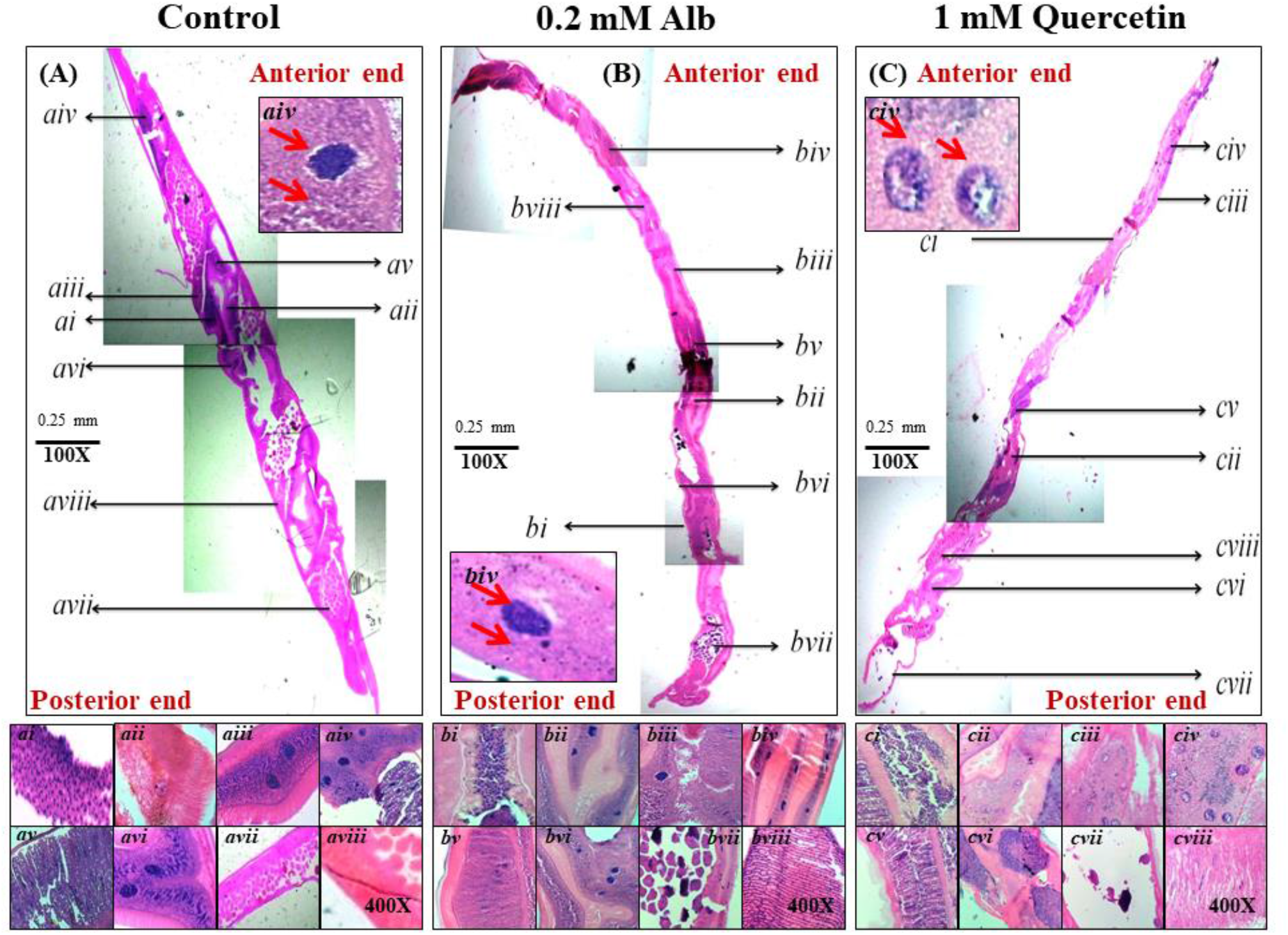
Histopathological changes in adult *H. contortus* after the treatment with quercetin. The first image (**a**) is showing the complete intact anterior and posterior ends along with other morphological features in the control worm, viz. isthmus, brut, pseudocoele, globular leukocytes (surrounded with nuclei), muscle cells, intestinal epithelial region, ovum, and growth zone of the ovary and intact skin tissue. Partial disorganizations were visible in some of these body parts after the treatment with Alb (**b**). After the treatment with a 1 mM of quercetin, completely disrupted anterior end together with other distorted areas, such as ovary, growth zone, pseudocoele, muscle cells, and wrecked globular leukocytes (with very less count of surrounding nuclei) were found (**c**). Due to the toxicity-mediated by quercetin, no eggs were found in the ovum (***cvii***). ***ai-aviii*** is showing isthmus, brut, pseudocoele, globular leukocytes, muscle cells, intestinal epithelial region, ovum and growth zone of the ovary, and intact skin tissue in control adult worm respectively. ***bi-bviii*** and ***ci-cviii*** are showing the same for Alb (0.2 mM) and quercetin (0.1 mM) treated worms respectively.

### 3.6. Generation of reactive oxygen species in the nervous system due to quercetin treatment

We investigated the mechanism underlying the toxicity elicited by quercetin and checked whether the compound is producing oxidative stress in the adult *H. contortus*. Reactive oxygen species (ROS) was found to be generated in the adult worm when treated with quercetin (1 mM) for 3 h (Fig. 3c, 3f, 3i, and 3l), whereas ROS was not detected in the control (Fig. 3a, 3d, 3g, and 3j) and Alb (Fig. 3b, 3e, 3h, and 3k) groups. Fig. 3c, 3f, 3i, and 3l are respectively showing the generations of ROS, in the adult worms, induced by quercetin in the anterior part of the body, ventral cord and tail ganglia (at the posterior end), nerve ring (at the anterior end), and commissural connections (middle of body region). No ROS was detected over the levels of autofluorescence at the same locations in the other two experimental groups. We also measured the increase in ROS generation in the nerve ring of the adult worm, caused by the treatment with 1 mM quercetin for 3 h, by repeated-measures ANOVA at different time points (after 1, 3, 5, and 7 min of DCFDA-treatment) and found a significant increase in pixel intensity (indicating increased staining with the ROS-detecting dye, DCFDA) with time (Fig. S4). Results of the repeated measures ANOVA showed a significant pixel-intensity × time effect (F_(60, 1600)_ = 1.67, *p* = 0.001) and a significant time effect (F_(3, 80)_ = 875.69, *p*<0.0001). Tukey’s HSD posthoc test revealed significant difference in pixel intensities between the following pairs of time points: 1 min vs. 5 min (*p* = 0.0001), 1 min vs. 7 min (*p* = 0.0001), 3 min vs. 5 min (*p* = 0.0001), 3 min vs. 7 min (*p* = 0.0001) 5 min vs. 7 min (*p* = 0.0001).

**Fig. 3:**
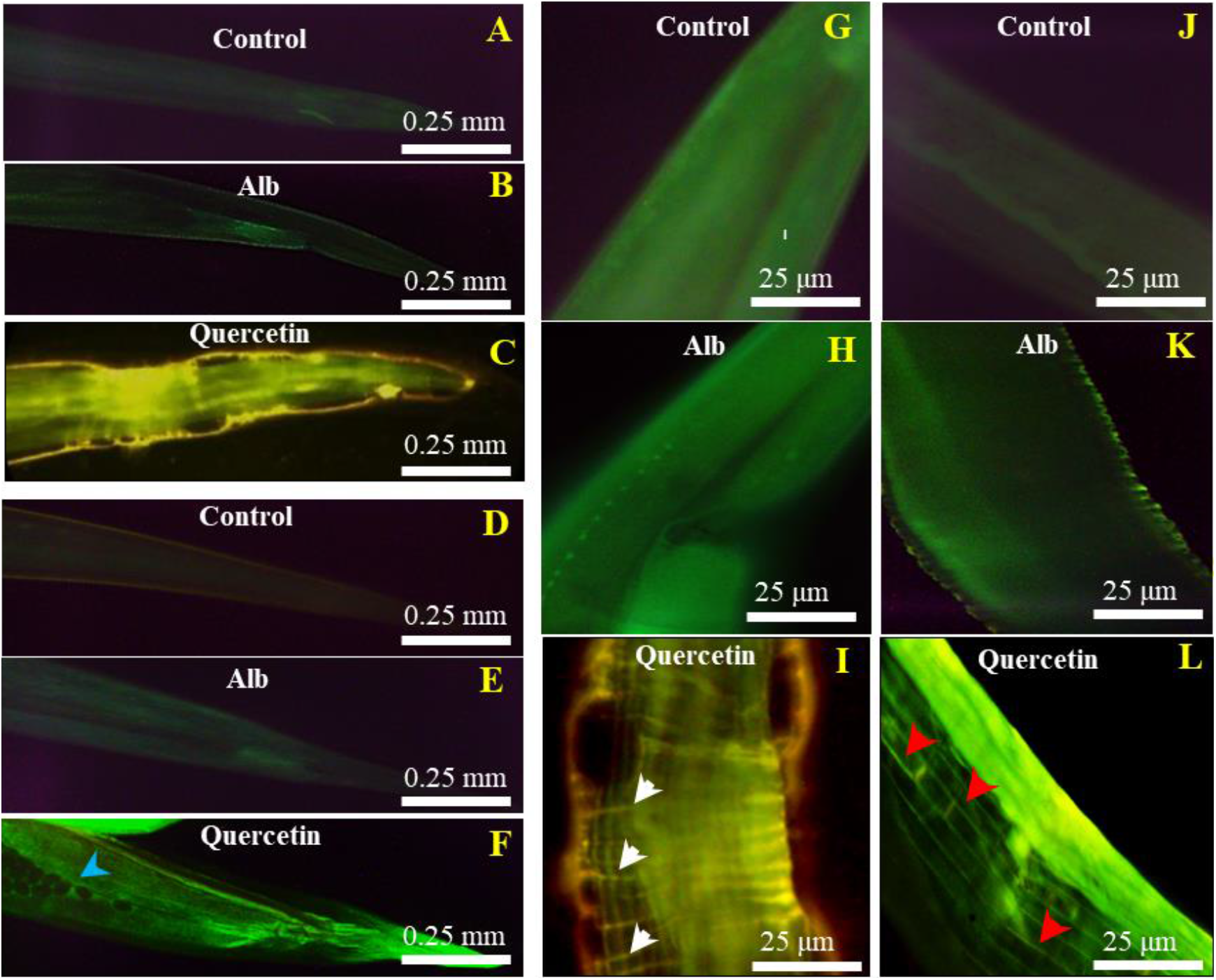
Generation of reactive oxidative species (ROS) in the nervous system of the adult worms due to exposure to quercetin. Adult worms were treated for 3 h with albendazole and quercetin and processed with the stain, DCFDA (100 nM) to detect the ROS. The images, from **a-c**, are showing the differences in staining (100 × magnification), in the anterior end, after 7 min of DCFDA-treatment (due to differences in ROS generation) respectively in the control (treated with RPMI media), Alb (0.2 mM) and quercetin (1 mM)-treated worms. The next three images (**d-f**) are showing the differences in staining (100 × magnification), in the posterior end, in the same three experimental groups. In the (**f**) the blue arrowhead is indicating the eggs of the quercetin-treated worm. Generations of ROS respectively in the nerve ring (**i)**, marked with white arrowhead; (400×), commissural connections **(l)**, marked with red arrowheads; (400×) and ventral cord, marked with yellow arrowhead; (400×) were detected in the adult worms treated with quercetin. Whereas no such structural discrimination and elevation of ROS was observed in control (**g, h**) and Alb treated (**j, k**) worms.

### 3.7. Alterations in the activities of catalase, superoxide dismutase, and glutathione peroxidase enzymes after quercetin treatment

Next, we have tested whether high ROS generation due to quercetin-mediated oxidative stress within the worm’s body has altered the activity levels of the enzymes involved in reducing oxidative stress, such as catalase (CAT), superoxide dismutase (SOD), and glutathione peroxidase (GPx). An increase in the catalase activity along with the increase in the concentration of quercetin was observed. The activity levels of CAT calculated respectively were 5.2±0.5, 5.9±0.1, 7.2±0.52, 16.1±0.18, and 16.6±0.2 U/mg proteins at the concentrations of quercetin, 0.125, 0.25, 0.5, 1.0 and 2.0 mM after 3 h of treatment (Table 2). A one-way ANOVA test found an increase in the activity level of CAT depending on the concentration of quercetin; significant interaction between the CAT-activity and quercetin-concentration (F_(5, 12)_ = 88765, *p*< 0.0001). Results of Tukey’s HSD posthoc test, comparing between the pairs of CAT-activity values, are given in Table ST3. A significant increase in CAT-activity was also found in the Alb-treated worms compared to the control (t = -25.84, df = 4, *p* = 0.00001), whereas no difference was found between the worms treated with 0.2 mM Alb and 0.25 mM quercetin (t = -2.14, df = 4, *p* = 0.09). A similar dose-response effect was also found for superoxide dismutase (SOD) as we calculated the activity levels respectively 2.13±0.56, 3.09±0.47, 4.02±0.56, 6.68±0.21, and 7.61±0.34 U/mg protein for the same set of concentrations of quercetin after 3 h of treatment (Table 2). One way ANOVA found a significant SOD-activity × quercetin-concentration effect (F_(5, 12)_ = 1398.3, *p*< 0.0001). Results of Tukey’s HSD posthoc test, comparing between the pairs of SOD-activity values, are given in Table ST4. Significant increase in SOD-activity was found in the Alb-treatment compared to the control (t = -26.08, df = 4, *p* = 0.00001), however no difference in activity levels was found between the groups, 0.2 mM Alb and 0.25 mM quercetin (t = 0.95, df = 4, *p* = 0.39). We also found the increasing activity of glutathione peroxidase, with increasing concentration of quercetin. The activity levels of GPx were found to be 6.32± 0.78, 9.39±0.51, 15.05±0.31, 17.28±0.56, and 18.47± 0.2 U/mg protein respectively concentrations of quercetin, 0.125, 0.25, 0.5, 1.0 and 2.0 mM after 3 h of treatment (Table 2). A one-way ANOVA has confirmed the dose-response for quercetin; significant GPx-activity × quercetin-concentration effect (F_(5, 12)_ = 25181, *p*<0.0001) (Results of the Tukey’s HSD posthoc test, comparing between the pairs of GPx-activity values, are given in Table ST5). Increased activity was also found for Alb-treatment compared to the control (t = -23.25, df = 4, *p* = 0.00001) and unlike the previous two enzymes, for GPx, we found a significantly higher activity in the Alb-treated worms than in the worms treated with 0.25 mM quercetin (t = -3.6, df = 4, *p* = 0.02).

**Table 2:** Alteration of activity level of different oxidative stress responsive enzymes of *H. contortus* due to quercetin treatment.

| Experimental Groups | U/mg protein |  |  |
| --- | --- | --- | --- |
|  | CAT | SOD | GPx |
| **Control** | 8.85±0.46 | 1.65±0.11 | 19.9±1.75 |
| **0.2 mM Alb** | 134.48±8.4 | 24.89±1.53 | 319.58±22.25 |
| **0.125 mM** | 107.86±0.64 | 18.36±0.08 | 183.61±0.97 |
| **0.25 mM** | 123.97±1.11 | 26.40±2.26 | 272.76±2.56 |
| **0.50 mM** | 149.6±0.42 | 34.63±0.08 | 436.47±1.68 |
| **1 mM** | 332.26±0.66 | 57.49±0.3 | 503.47±0.48 |
| **2 mM** | 342.08±0.72 | 63.30±0.19 | 518.33±1.75 |
| **5 mM** | 363.93±0.42 | 65.47±0.23 | 536.27±1.28 |

## 4. Discussion

Anthelmintic resistance against the broad-spectrum anthelmintics, used for the gastro-intestinal nematodes, is becoming rampant due to their widespread use [77,78]. The severity of helminthic infection can be understood from the WHO reports that claimed death in tens of thousands and millions of disablements annually due to the intestinal helminthic infections among which infants and pregnant women are the most vulnerable groups, especially in countries with poor hygiene and economic status[79,80]. In the quest for alternative drugs against the parasitic nematodes, research efforts were invested to evaluate the effectiveness of plant metabolites, including the flavonoids[81].

The present study the anthelmintic potential of a previously known flavonoid, quercetin, in *H. contortus* isolates from the state of Punjab, India is found to exert a nematicidal effect in the concentration range from µM to mM and to be most effective at the concentration of 1 mM. At this concentration, adult male worms manifested 80 % paralysis after 3 h of treatment and 100 % were found dead after 24 h. Female, although showed more resistance to 1 mM quercetin than the males as the paralysis of 80 % of the adult females took 6 h, however, 100 % death of the females was observed at 24 h of exposure, like the males. Likewise, the LD_50_ value for the adult females was higher than that of the males for quercetin. The higher susceptibility of the males in comparison to the females could be due to the smaller size of the males than the females[82]. Alternatively, the genetic constitution of the females has probably contributed as it is reported that female *H. contortus* showed increased resistance to benzimidazole and thiabendazole, even if thiabendazole resistance is probably not a sex-linked trait [46,83]. Differential susceptibility among the sexes was also found in the evaluation of anthelmintic efficacy of the extracts of the plant, *Hedera helix*, against *H. contortus*, which showed a dose-dependent improvement for the male than the female parasites[55].

In the behavioral domain, we also found larval death and inhibition to the process of egg hatching due to quercetin treatment. Quercetin showed dose-dependent larval death and inhibition of egg hatching in our assays which confirmed the concentration-dependent lethal effects of the flavonoid on the different developmental stages of *H. contortus*. Substantial death of the larvae (nearly 90% death) and inhibition of egg hatching (nearly 80 % of the eggs fail to hatch) are found after the treatment with 1 mM concentration of quercetin respectively for 24 h and 48 h. Besides, quercetin showed higher potential than the popularly used anthelmintic, albendazole. Whereas albendazole, caused only 44 % death of the larvae and exerted 28 % inhibition in egg hatching at a concentration of 0.2 mM, quercetin at a similar concentration (0.25 mM) caused 67 % larval death and showed 65 % inhibition in egg hatching. The lethal dosages for the larval (24 h) and egg (48 h) stages calculated were 0.16 mM and 0.19 mM, respectively. These results together with the paralysis and death time analysis establish the increased efficacy of quercetin as an anthelmintic agent against the egg, larval and adult stages of *H. contortus*.

The toxic nature of quercetin has also been further proven by its detrimental effects on the physical and histological structures of the adult *H. contortus*. Physical damages inflicted on the adult worms, as monitored by the scanning electron microscopy, revealed that quercetin ruptured the cuticular structure of the body and completely disrupted the two ends and other body regions, including the destruction of the blood-sucking anterior mouth part and the posterior tail region. Physical damage, in the forms of partial shrinkage of the body, the body ends, and disruption of the cuticle is also visible for albendazole treatment, however, the extent of damage caused by quercetin is much more than albendazole. In our study, for the first time, we reported the histological constitutions of *H. contortus* and apoptotic histopathology, caused by quercetin. Using the technique of Hematoxylin & Erythrosin (HnE) staining, we identified the isthmus, brut, pseudocoele, globular leukocytes, muscle cells, intestinal epithelial region, and growth zone of the ovary in the different sections of the body of *H. contortus*. We demonstrated detrimental changes in the morphologies of these body regions when treated with quercetin as compared to the control and albendazole. Whereas, albendazole treatment, partly breached the muscle cells and ovary along with some disruptions were visible in the isthmus and pseudocoele, quercetin at 1 mM drastically alter these structures with a complete loss of eggs in the ovary and damaged globular leukocytes.

Irrespective of the promising lethal effect of quercetin on *H. contortus*, the potential use of this compound as an anthelmintic drug demands a clear understanding of its mechanism of action. We thus further investigated the toxicity mechanism of quercetin in the adult *H. contortus* and found that the compound brought about oxidative stress in this parasitic helminth, targeting the nervous system. Deformations of the body parts (body area and the anterior and posterior body ends), due to exposure to quercetin for 3 h, were found to be associated with the generation of the high amount of reactive oxygen species (ROS) in the nerve ring, ventral cord and commissural connections in the adult worms. Besides, we found that quercetin treatment leads to the production of a higher amount of ROS than treatment with albendazole as well as a significant increase in ROS generation with time was seen in the nerve ring area due to quercetin exposure. This burden of ROS-induced oxidative stress in the nervous system may be the cause of paralysis and eventual death of the adult worms because oxidative stress in the nervous system has been reported in numerous studies to cause adverse and irreversible modifications of the cellular macromolecular machinery and eventual death of the cell, including neurodegeneration through multiple mechanisms[84–86].

Cell however respond to the ROS-induced oxidative stress by upregulating the activities of enzymes that comprise the antioxidant defense system of the cell, by scavenging the ROS (like superoxide radicle and hydrogen peroxide), such as superoxide dismutase (SOD), catalase (CAT), glutathione peroxidase (GPx), thioredoxin peroxidase, glutathione S-transferase[87]. In our *in vitro* assays, we estimated the activity levels of three important enzymes viz CAT, SOD, and GPx in the tissue homogenate of adult *H. contortus* after 3 h of treatment with quercetin and Alb. We found quercetin-mediated dose-dependent responses for all three enzymes which correlate well with the fluorescence imaging data, confirming a high amount of ROS generation in the nervous system of the parasitic helminth, *H. contortus*, as increasing concentration of quercetin leads to significant elevations of activities of these antioxidant enzymes. Alb treatment also showed comparable activity levels of these enzymes with quercetin, except for GPx, in which Alb-treatment resulted in higher activity than the comparable dose of quercetin. Thus the change in metabolism, as evident from the upregulations of antioxidant enzyme activities, mediated by quercetin is the biochemical signature of its lethal effect which contributes to the physical damage, histopathology, paralysis, and eventual death of this pathogenic nematode. Altogether, our study, for the first time, confirmed the nematicidal effects of quercetin with 1 mM concentration found to be effective against the egg, larval, and adult stages of *H. contortus*. This is also the first report of histopathology in *H. contortus* worms. Moreover, the results demonstrated the higher antihelminthic potential of quercetin than the popular drug, albendazole, which is extremely significant from the perspective of increasing resistance of *H. contortus*, worldwide, against the traditional antihelminthic drugs. Although we couldn’t test whether our samples of *H. contortus* had any prior resistance against albendazole, however the mechanism of action of quercetin opens the window of further research to establish a novel antihelminthic remedy. The *in vitro* studies conducted with quercetin are extremely useful for screening the efficacy of any potential anthelmintic agent against the worms, especially which are resistant to the commercially available drugs. Furthermore, the study also emphasizes the significance of using quercetin in the *in vivo* models for treatments against the gastrointestinal parasitic helminths. However, our current understanding of the functional mechanisms of the nervous system of parasitic helminths, from physiology to molecular biology is very limited. The same also applies to pharmacology owing to the lack of understanding of the nature of anthelmintics as whether they are cell-type specific toxins or metabolic toxins or disrupting other pathways, as well as our poor knowledge on the dynamics of the drugs inside different locations of the nematodes. These two factors put serious constraints to conclude on the functioning of anthelmintic drugs, especially when a drug proves to be effective at higher concentrations (in the millimolar range), like in our assays we found quercetin to be highly effective at 1 mM concentration. We thus believe that quercetin at this high concentration exerted adverse effects not only on the nervous system but also on other tissues which are also visible in the substantial increase in the activity levels of the antioxidant enzymes collected from the whole body of the adult *H. contortus*. Additionally, the lethal effect of quercetin on the nervous system of *H. contortus* is contrary to the findings in mammalian systems, where quercetin acts as a neuroprotective agent. We speculate that the same drug may not work in the same way in the nematodes and mammals because the repertoires of the receptor and the downstream signaling molecules may not be the same amongst them. Thus the case for intensive research in this direction is strong which will decipher the detailed molecular mechanisms of pharmacology in the parasitic helminths.

## CONCLUSION

This is the first report of histopathology in *H. contortus* worms mediated by quercetin, a plant flavonoid that is present ample in nature. We showed that quercetin has prominent nematicidal effects at 1 mM concentration against the egg, larval and adult stages of *H. contortus*. The results also demonstrated the higher antihelminthic potential of quercetin than the popular drug, albendazole. We found that quercetin induces oxidative stress in the adult stage of the worm, prominently in their nervous system, which causes a predominant change in metabolism and is probably associated with physical damage, and eventual death of the worms.

## ACKNOWLEDGMENTS

DC is thankful to TIET-VT-CEEMS (CEEMS-TOF/2020) for funding the lab and NKC is thankful for the seed money grant, TSLAS, TIET. VG and SS are thankful to TIET for fellowship. V.G. and S.S. is thankful to TIET for fellowship.

## AUTHOR CONTRIBUTIONS

Conceptualization: V.G, D.C; Methodology development: V.G, L.D.S., D.C.; Data Collection: V.G., S.S.; Data analysis: V.G., N.K.C; Supervision: D.C.; Visualization: V.G., N.K.C., S.S; Writing original draft: V.G.; Writing – review & editing: D.C., N.K.C, L.D.S.; Funding acquisition: D.C.

