## Supplementary Figures and Tables for "Targeting the nervous system of the parasitic worm, *Haemonchus contortus*, with quercetin"

.

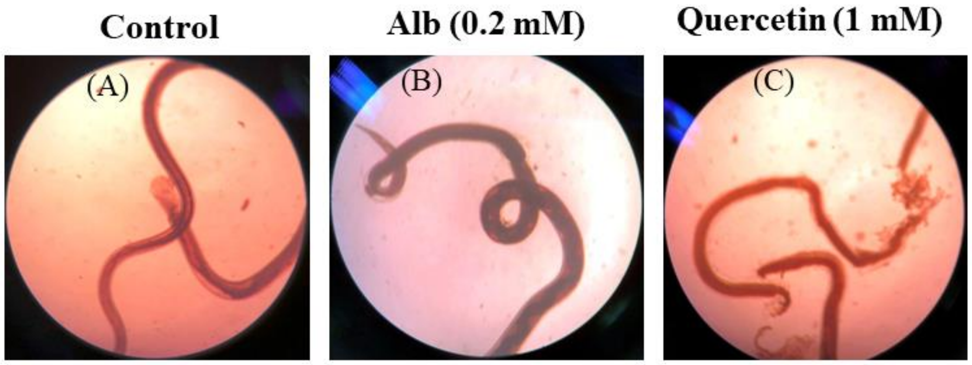


**Fig. S1: Physical damage of the adult female *H. contortus* due to quercetin exposure.** Bright-field microscopy revealed physical damage in the adult female worms (A-C) due to exposure to quercetin for 24 h. Quercetin inflicted more physical damage than albendazole, whereas the control condition didn't cause any damage to the worms.

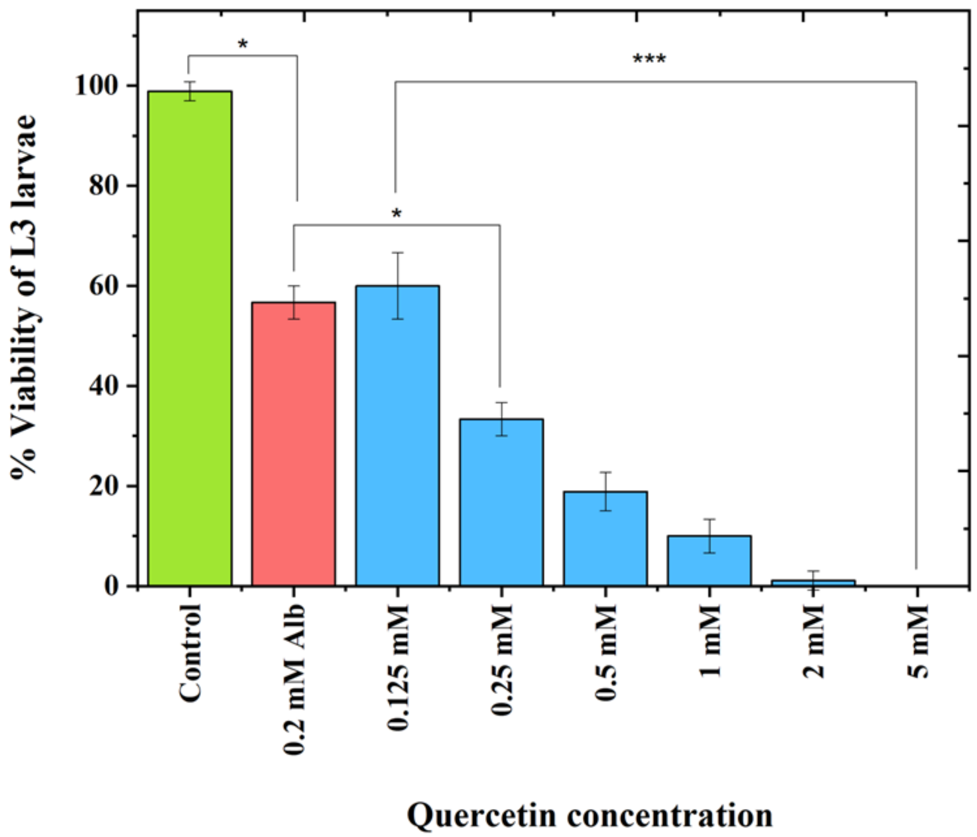


**Fig. S2: Percentviability of L-3 larval stage of *H. contortus* after treatments with different concentrations of quercetin and Alb.** The blue bars are showing the percent viability of the L3 larval stage after 24 h of treatment different concentrations of quercetin (0.125, 0.25, 0.50, 1, 2, and 5 mM). The green and red-colored bars are respectively representing the survival of larvae under the control condition (RPMI media) and after albendazole (0.2 mM) treatment. Quercetin treatment showed concentration-dependent effects on the survival of the L3 larvae and higher mortality than Alb. The results of the three experiments are plotted as mean±SD.

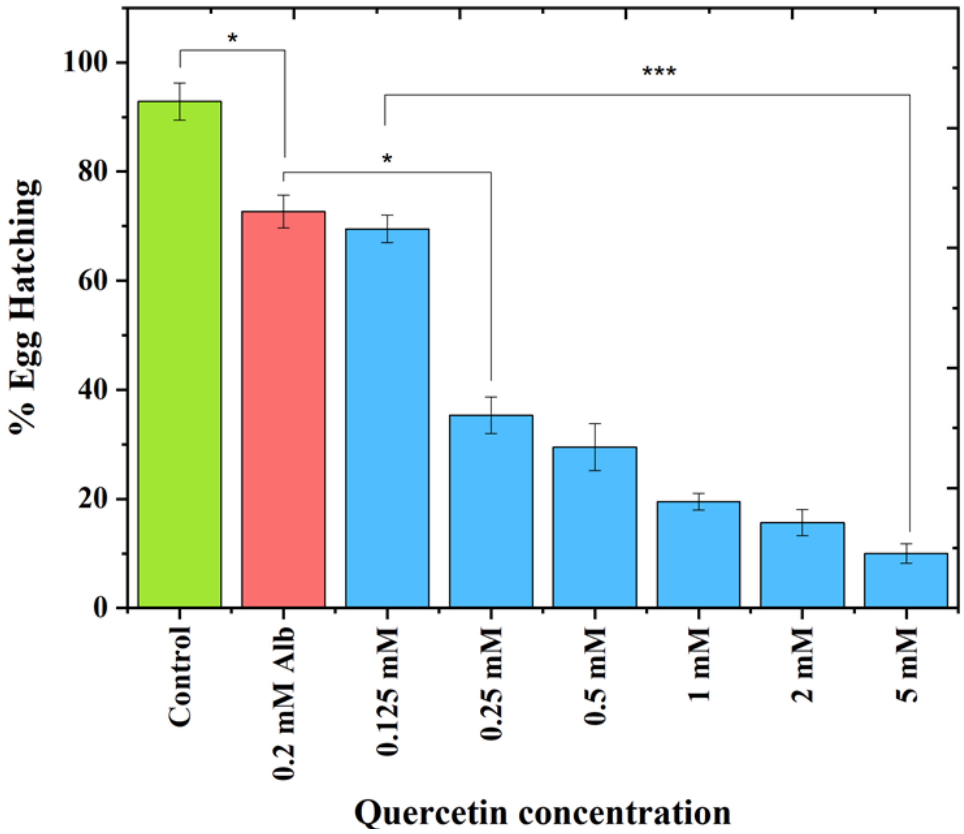


**Fig. S3: Percentage of eggs hatched after treatments with different concentrations of quercetin and Alb.** Percentage of eggs, hatched and released L1 larvae, were calculated and plotted as bar graphs after 48 h treatment with RPMI media (green bar: control condition), different concentrations of quercetin (blue bars: 0.125, 0.25, 0.5, 1, 2, and 5 mM) and albendazole (red bar: 0.2 mM). The results of three independent experiments are presented as means with the corresponding standard deviations. Quercetin treatment showed a concentration-dependent change in the hatching of eggs and lower hatching than Alb.

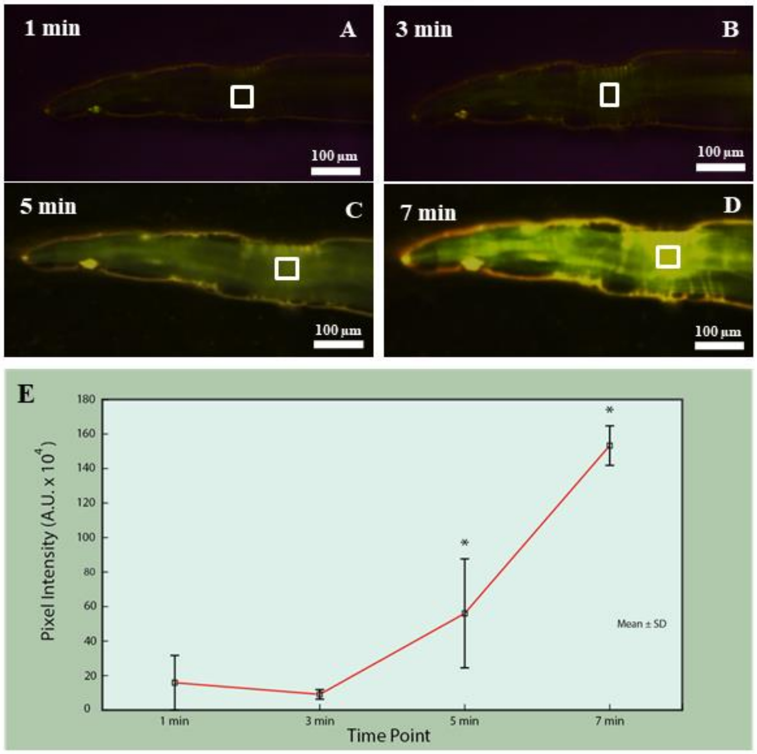


**Fig.S4: Increase in DCFDA staining in the nerve ring area of adult worm with time, after quercetin treatment.** DCFDA treatment of the adult worm, after 3 h of quercetin treatment, showed a significant increase in staining in the nerve ring area with time. An increase in staining, represented by an increase in the average pixel intensity, was due to higher amounts of ROS generation with time as a result of quercetin treatment. Repeated measures ANOVA showed a significant pixel-intensity × time interaction effect ($F_{(60,\ 1600)}$= 1.67, $p$= 0.001) as well as a significant time effect ($F_{(3,\ 80)}$ = 875.69, $p$<0.0001).

**Table S1:** Results of Tukey's HSD posthoc test that follows the one-way ANOVA are shown here. Survival of L3 larvae for the pairs of quercetin concentration is compared. Differences are considered statistically significant if the corresponding p-value was <0.01. P-values that are not significant are denoted in bold.

| | 0.125 mM | 0.25 mM | 0.5 mM | 1 mM | 2 mM | 5 mM |
|---|---|---|---|---|---|---|
| 0.125 mM | | | | | | |
| 0.25 mM | 0.0001 | | | | | |
| 0.5 mM | 0.0001 | 0.005 | | | | |
| 1 mM | 0.0001 | 0.0002 | **0.1** | | | |
| 2 mM | 0.0001 | 0.0001 | 0.001 | **0.1** | | |
| 5 mM | 0.0001 | 0.0001 | 0.0006 | 0.06 | **0.9** | |

**Table S2:** Results of Tukey's HSD posthoc test that follows the one-way ANOVA are shown here. The hatching of eggs for the pairs of quercetin concentration is compared. Differences are considered statistically significant if the corresponding p-value was <0.01. P-values that are not significant are denoted in bold.

| | 0.125 mM | 0.25 mM | 0.5 mM | 1 mM | 2 mM | 5 mM |
|---|---|---|---|---|---|---|
| 0.125 mM | | | | | | |
| 0.25 mM | 0.0001 | | | | | |
| 0.5 mM | 0.0001 | 0.1 | | | | |
| 1 mM | 0.0001 | 0.0003 | 0.008 | | | |
| 2 mM | 0.0001 | 0.0001 | 0.0007 | **0.5** | | |
| 5 mM | 0.0001 | 0.0001 | 0.0001 | 0.01 | **0.2** | |

**Table S3:** Results of Tukey's HSD posthoc test comparing the activity levels of catalase for the pairs of quercetin concentration. Differences are considered statistically significant if the corresponding p-value was <0.01.

| | 0.125 mM | 0.25 mM | 0.5 mM | 1 mM | 2 mM | 5 mM |
|---|---|---|---|---|---|---|
| 0.125 mM | | | | | | |
| 0.25 mM | 0.0001 | | | | | |
| 0.5 mM | 0.0001 | 0.0001 | | | | |
| 1 mM | 0.0001 | 0.0001 | 0.0001 | | | |
| 2 mM | 0.0001 | 0.0001 | 0.0001 | 0.0001 | | |
| 5 mM | 0.0001 | 0.0001 | 0.0001 | 0.0001 | 0.0001 | |

**Table S4:** Results of Tukey's HSD posthoc test comparing the activity levels of superoxide dismutase for the pairs of quercetin concentration. Differences are considered statistically significant if the corresponding p-value was <0.01. P-values that are not significant are denoted in bold.

| | 0.125 mM | 0.25 mM | 0.5 mM | 1 mM | 2 mM | 5 mM |
|---|---|---|---|---|---|---|
| 0.125 mM | | | | | | |
| 0.25 mM | 0.0001 | | | | | |
| 0.5 mM | 0.0001 | 0.0001 | | | | |
| 1 mM | 0.0001 | 0.0001 | 0.0001 | | | |
| 2 mM | 0.0001 | 0.0001 | 0.0001 | 0.0002 | | |
| 5 mM | 0.0001 | 0.0001 | 0.0001 | 0.0001 | **0.12** | |

**Table S5:** Results of Tukey's HSD posthoc test comparing the activity levels of glutathione peroxidase for the pairs of quercetin concentration. Differences are considered statistically significant if the corresponding p-value was <0.01.

| | 0.125 mM | 0.25 mM | 0.5 mM | 1 mM | 2 mM | 5 mM |
|---|---|---|---|---|---|---|
| 0.125 mM | | | | | | |
| 0.25 mM | 0.0001 | | | | | |
| 0.5 mM | 0.0001 | 0.0001 | | | | |
| 1 mM | 0.0001 | 0.0001 | 0.0001 | | | |
| 2 mM | 0.0001 | 0.0001 | 0.0001 | 0.0001 | | |
| 5 mM | 0.0001 | 0.0001 | 0.0001 | 0.0001 | 0.0001 | |
